# Differential Role of GABAergic and Cholinergic Ventral Pallidal Neurons in Behavioral Despair, Conditioned Fear Memory and Active Coping

**DOI:** 10.1101/2022.07.21.500949

**Authors:** Cemal Akmese, Cem Sevinc, Sahar Halim, Gunes Unal

**Author notes:** Equal contribution (Cemal Akmese, Cem Sevinc).

## Abstract

The ventral pallidum (VP), a core structure of the reward circuit, is well-associated with appetitive behaviors. Recent evidence suggests that this basal forebrain nucleus may have an overarching role in affective processing, including behavioral responses to aversive stimuli. We investigated this possibility by utilizing selective immunotoxin lesions and a series of behavioral tests in adult male Wistar rats. We made bilateral GAT1-Saporin, 192-IgG-Saporin or PBS (vehicle) injections into the VP to respectively eliminate GABAergic and cholinergic neurons, and tested the animals in the forced swim test (FST), open field test (OFT), elevated plus maze (EPM), Morris water maze (MWM) and cued fear conditioning. Both GAT1-Saporin and 192-IgG-Saporin injections reduced behavioral despair without altering general locomotor activity. This antidepressant-like effect was accompanied by reduced freezing and increased darting in the acquisition phase of cued fear conditioning. In the test phase, cholinergic, but not GABAergic lesions, impaired context-dependent fear memory, while both groups showed diminished conditioned freezing in a novel context. In line with this, selective cholinergic lesions impaired spatial memory in the MWM as compared to GAT1-Saporin or vehicle-injected animals. No differential effect was observed in anxiety-like behavior assessed in the OFT or EPM. These findings show that both the GABAergic and cholinergic cell groups of the VP contribute to behavioral despair and acquired fear responses by suppressing active coping. In conditioned learning, cholinergic ventral pallidal neurons contribute to the fear response in a context-independent manner, while the GABAergic population is required when the context information is missing.

**Significance Statement:** The ventral pallidum (VP), associated with motivation and appetitive behaviors, is tested for its role in behavioral responses to aversive stimuli. Findings show that both the GABAergic and cholinergic neuronal groups of the VP contribute to behavioral despair and acquired fear response by suppressing active coping. In conditioned learning, the cholinergic neurons are required for the acquired fear response, while the VP GABAergic population becomes necessary only when the context information is missing. These results suggest that ventral pallidal GABAergic and cholinergic projections to the amygdaloid complex may constitute therapeutic targets for affective disorders.

## Introduction

The ventral pallidum (VP) is a basal forebrain nucleus acting as a functional interface for translating motivation into motor action (Lengyel et al., 2005). The VP, possessing reciprocal connections with the nucleus accumbens (NAc) and the ventral tegmental area (VTA) (Groenewegen et al., 1993; Haber et al., 1985; Hakan et al., 1992; Heimer et al., 1991; Phillipson & Griffiths, 1985; Root et al., 2015; Taylor et al., 2014; Zahm et al., 1996), has been conceptualized as an output station of the dopaminergic mesolimbic pathway (Haber et al., 1985; Pierce & Kumaresan, 2006; Smith et al., 2009) and a core part of the “reward circuit” of the brain (Haber & Knutson, 2010). Behavioral data clearly show the role of VP in reward processing (Ottenheimer et al., 2020; Richard et al., 2016; Smith et al., 2009; Stephenson-Jones, 2019) and hedonic liking (Ho & Berridge, 2013; Smith & Berridge, 2005). In addition to its role in processing appetitive stimuli, the VP is also involved in responses to aversive stimuli (Wulff et al., 2019). Non-specific VP lesions lead to enhanced food aversion (Cromwell & Berridge, 1993), while recent evidence suggests that VP neurons may encode different types of aversive information (Farrell et al., 2021; Moaddab et al., 2021; Saga et al., 2017; Stephenson-Jones et al., 2020). Disturbing VP glutamatergic activity alters behavioral avoidance (Faget et al., 2018; Stephenson-Jones et al., 2020) and inhibition of GABAergic activity leads to heightened risk aversion (Farrell et al., 2021). Moreover, VP neuronal subpopulations may contribute to distinct rodent phenotypes of clinical depression (Knowland et al., 2017). Utilizing a comprehensive experimental design based on the emerging evidence, this study attempts to elucidate specific functions of the VP in behavioral responses to aversive stimuli. We selectively lesioned GABAergic and cholinergic neurons of the VP, and tested the animals for behavioral despair, locomotor activity, anxiety, spatial learning and conditioned fear memory.

The VP contains co-distributed populations of GABAergic and cholinergic neurons (Gritti et al., 2006; Gritti et al., 1993) that are reciprocally connected with the amygdaloid complex (Agostinelli et al., 2019; Carlsen et al., 1985; Do et al., 2016; Hintiryan et al., 2021; Mascagni & McDonald, 2009; Mcdonald et al., 2011a; Záborszky et al., 1984). Similar to the GABAergic septo-hippocampal projections (Freund & Antal, 1988; Unal et al., 2015), approximately 80-90% of amygdala-projecting VP GABAergic neurons selectively synapse onto GABAergic interneurons in the amygdala (Mcdonald et al., 2011a). This suggests that a similar spatio-temporal inhibition mechanism imposed by GABAergic septo-hippocampal projections (Tóth et al., 1997; Unal et al., 2015) may exist for VP GABAergic neurons targeting the amygdala. Recent functional evidence shows that the VP is involved in amygdala-dependent limbic processes in addition to its central position in the mesolimbic/reward pathway, suggesting a role for this basal forebrain nucleus in the integration of positive and negative valence. Encoding and differentiating appetitive and aversive stimuli are crucial for adaptive behaviors, and dysfunction in these processes is associated with a wide range of mood disorders (Nusslock & Alloy, 2017). However, the specific functions of distinct VP neuronal subpopulations in aversive conditions and their relation to appetitive behaviors are yet to be characterized.

In the current study, we used saporin-based immunotoxins to selectively lesion the GABAergic (GAT1-Saporin) or cholinergic (192-IgG-Saporin) neurons in the VP (Bolshakov et al., 2020). We performed a battery of behavioral tests to assess alterations in behavioral despair, locomotor activity, anxiety-like behavior, hippocampus-dependent spatial learning, active behavioral coping and conditioned fear responses. We found that both GAT1-Saporin and 192-IgG-Saporin injections reduced behavioral despair without altering general locomotor activity. Both groups displayed reduced freezing and enhanced active coping during cued fear conditioning, while GABAergic and cholinergic VP neurons differentially contributed to extinction in a context-dependent manner. Similarly, cholinergic, but not GABAergic VP lesions impaired performance in hippocampus-dependent spatial learning.

## Materials and Methods

### Animals

Twenty-four adult male Wistar rats (240-320 g) were used for the experiments. They were housed in groups of four under standard vivarium conditions (21 ± 1 °C; ~ 50% humidity; 12:12 day/night cycle with lights on at 8:00 AM) with *ad libitum* access to food and water prior to stereotaxic surgery. They were then transferred to individual cages and remained there until the end of the experiment. All procedures were approved by the Boğaziçi University Ethics Committee for the Experimental Use of Animals in Scientific Research.

### Stereotaxic Surgery

Animals were deeply anesthetized with an IP injection of a mixture of ketamine (85 mg/kg) and xylazine (8.5 mg/kg). Once reflexes were lost, we shaved their heads and secured them into a stereotaxic frame (Kopf Instruments). Body temperature was kept at approximately 37 °C during the surgery with a heating pad. A local anesthetic (Vemcaine, 10% lidocaine) was applied on the head and a vertical incision was performed to expose the skull. Injection coordinates of the VP (AP = −0.20, ML = ±2.20, DV = −7.60) were determined with reference to the Bregma point according to the rat brain atlas of Paxinos and Watson (2007). Following craniotomy, each animal received bilateral Gat1-Saporin (Advanced Targeting Systems, 325 ng/μl in Phosphate Buffered Saline (PBS), 0.5 μl volume, 0.1 μl/min), 192-IgG-Saporin (Advanced Targeting Systems, 500 ng/μl in PBS, 0.5 μl volume, 0.1 μl/min), or vehicle (PBS, 0.5 μl volume, 0.1μl/min) injections via 1 μl-Hamilton syringes attached to a microinjection pump. In order to minimize dorsal diffusion, each syringe was left in the injection site for five minutes at the end of the injection. The incision was sutured and local anesthetics (Anestol pomade, 5% lidocaine and Jetokain, 5 mg/kg, SC) and an antiseptic (Batticon) were applied to the stitch.

At the end of the surgery, the animals were taken to the post-operative care unit, placed under an infrared lamp and monitored until the effects of anesthesia wore off. Each animal went through a 14-day post-surgery recovery period to ensure complete healing and maximize immunotoxin-based elimination of the targeted cellular population before behavioral testing. To facilitate detection of somatic GABA in immunohistochemistry, four animals from the GAT1-Saporin and vehicle groups underwent a second stereotaxic surgery two weeks after behavioral testing and received bilateral colchicine injections into the lateral ventricles. The same procedures were repeated in these surgeries. Intracerebroventricular injections of colchicine (50 μg/5 μl in 0.9% saline for each hemisphere) were performed, instead of immunotoxin or PBS injections using 10 μl-Hamilton syringes.

### Experimental Design

Behavioral testing started with the 15-minute-long pretest or acclimation session of the forced swim test (FST) as visualized in Figure 1. Following a 24-hour break, each animal went through the 5-minute test session of the FST. Animals were tested for general locomotor activity in an open field test (OFT) on Day 3 and anxiety-like behavior in an elevated plus maze (EPM) on Day 4. A 5-day Morris water maze (MWM) protocol started on Day 5 and ended with the probe trial on Day 9. The final behavioral test was the auditory fear conditioning conducted on Day 10. Two extinction sessions were run on Days 13 and 17, first in the same apparatus, and then in a novel context. In OFT, EPM and fear conditioning, mazes/chambers were cleaned with 70% ethanol and dried using paper towels between sessions to eliminate olfactory cues for the next animal. Bilateral intracerebroventricular colchicine injections were made (Vehicle: n = 2, GAT1-Saporin: n = 2) 2 weeks after behavioral testing. Animals underwent perfusion-fixation following behavioral testing (n = 20) or colchicine injections (n = 4).

**Figure 1.**
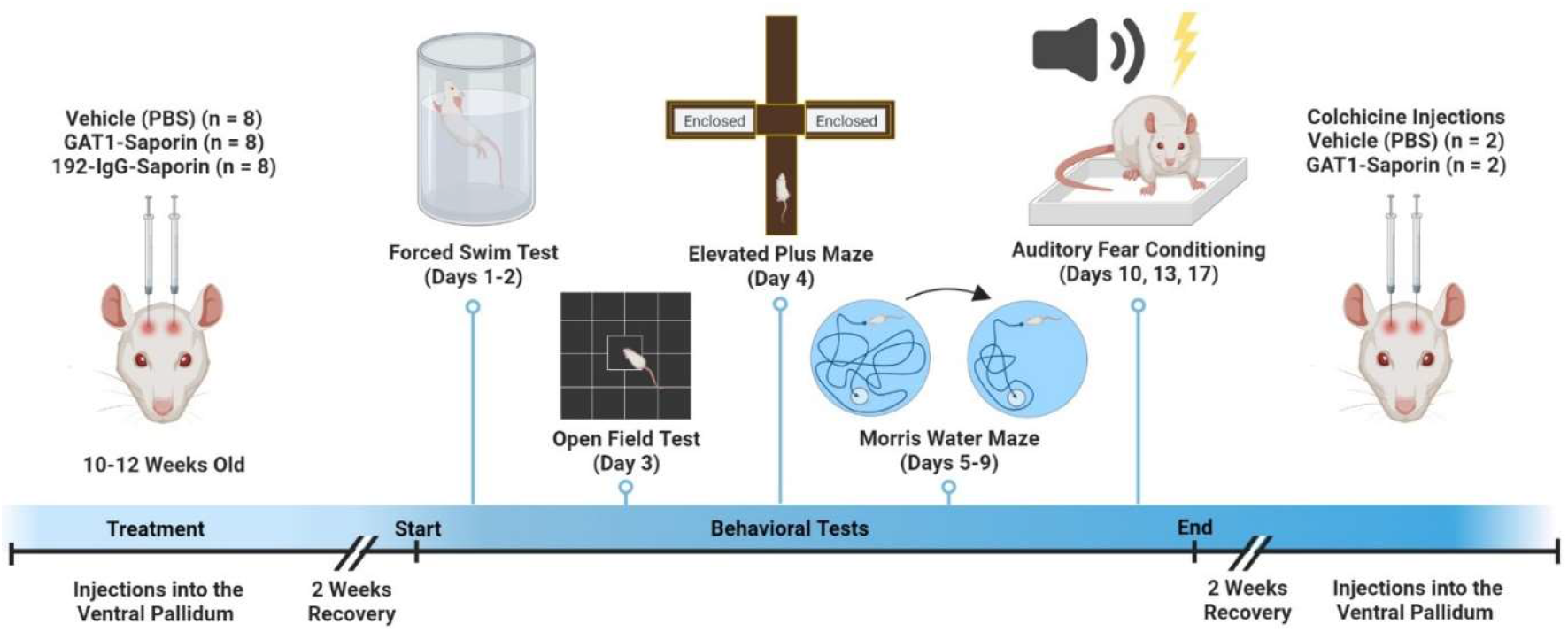
Experimental timeline illustrating the order of stereotaxic surgeries, recovery periods and behavioral testing. Figure created with BioRender.com.

### Forced Swim Test (FST)

We followed the standard FST protocol for rats (Porsolt et al., 1978). Animals were placed into a water-filled cylinder (depth = 30 cm, temperature = 25 ± 1 °C) and tested twice with a 24-hour break between the pretest (15 min) and test (5 min) sessions. Before each trial, the animal was brought to the behavioral testing room and was allowed to get acclimated to the environment for 5 minutes. At the end of each trial, the animal was dried with a paper towel, remained under an infrared heating lamp for 30 minutes and taken back to its home cage.

Behavioral analyses of the test session included recording overall time of immobility, swimming and climbing behavior with a time-sampling method (Slattery & Cryan, 2012). The 5-minute session was virtually divided into 5-second bins and the predominant behavior in each bin was recorded. Climbing behavior was defined as any active upward-directed motion where the animals’ paws were above the water surface whereas swimming was defined as any active movements parallel to the water surface. We also recorded the number of headshakes and dives, two common active behaviors in the FST. Bins in which diving or headshaking were the predominant behavior were recorded as climbing behavior. Each animal was coded by two observers who were blind to the experimental conditions, and the arithmetic mean of their scores were used for statistical analyses.

### Open Field Test (OFT)

Each animal was given a 5-minute acclimation period in the test room before the OFT. Animals, then, were placed in an opaque black colored open field (70 × 70 × 45 cm) for 5 minutes. The time spent in the central area (1600 cm^2^) versus periphery (within 15 cm of maze walls), overall locomotion (the average speed and total distance traveled) and the number of supported and unsupported rearing behaviors were recorded. Sessions were taped with a video camera and behavioral analyses were performed with a modified version of DeepLabCut (DLC) (Mathis et al., 2018) implementation.

### Elevated Plus Maze (EPM)

Anxiety-like behavior was assessed with an EPM positioned 50 cm above the ground (Pellow et al., 1985). It consisted of 2 acrylic transparent (open) and 2 wooden opaque (closed) arms (each arm: 50 × 10 cm). Animals were placed at the center of the maze facing one of the open arms. Each session lasted 5 minutes. The light intensity in different arms were recorded before the session to ensure that illuminance was significantly different in open (120 lx) and closed arms (30 lx). Overall time spent in the open versus closed arms and the number of crossings between the arms were recorded using DLC.

### Morris Water Maze (MWM)

We used a standard MWM (diameter = 120 cm, water temperature: 21 ± 1°C) to assess hippocampus-dependent spatial learning and memory. The protocol consisted of 4 days of training followed by a single probe trial (Vorhees & Williams, 2006). A circular escape platform (diameter = 10 cm) was placed 2 cm below the water level until it was removed for the probe trial. The water in the maze was opacified with milk powder in order to visually obscure the location of the escape platform. Four distinct proximal cues, 90° apart, were attached to the walls of the plexiglass maze.

Each training day consisted of four consecutive trials. Animals began each trial from a different starting point. Animals were allowed to stay on the platform for 15 seconds before being removed from the maze. The maximum amount of time allowed to locate the hidden platform was 2 minutes. If an animal could not find the platform during this time, it was gently led to the platform by its tail and left there for 15 seconds. Latency to locate the platform (escape latency), average swimming speed, duration spent in each quarter and the total distance swam were recorded and analyzed in the DLC.

On the probe trial day, the platform was removed from the maze and the animals were tested in a single session for 60 seconds. We assessed the time spent in the target quadrant where the platform was located, number of crossings over the original place of the platform, total distance travelled, average speed and thigmotaxis.

### Fear Conditioning

We utilized Pavlovian fear conditioning by pairing an auditory conditioned stimulus (CS; 80 dB, 2kHz, 6 sec) with a mild footshock (US; 1.0 mA, 2 sec). Delay conditioning carried out on Experimental Day 10 was followed by extinction sessions in the same conditioning chamber (context A) on Day 13 and in a different box (context B) on Day 17 (Fig. 6A).

The fear conditioning apparatus (context A) was a 25 × 40 × 20 cm chamber with transparent dark grey acrylic walls. Its floor consisted of 16 metal bars, each 2.5 cm apart, connected to a custom-made stimulator via an Arduino microcontroller. Following a 3-minute baseline measurement phase in the chamber, five CS-US pairings were presented with an intertrial interval (ITI) of 66 seconds. The extinction sessions consisted of a 3-minute-long baseline measurement followed by 12 presentations of the CS (ITI = 66 sec). The second extinction session (Context B) was conducted in a 35 × 35 cm, open top square chamber with a black solid floor, one transparent and three 3 opaque walls. It was also located in a different part of the behavioral testing room.

All sessions were recorded by two video cameras, one positioned above and the other in front of the conditioning/extinction chambers. Duration of freezing to US or CS presentations, and the number of darting and jumping responses to the US were coded offline by two observers who were blind to the experimental conditions.

### Histology and Immunohistochemistry

Perfusion-fixations were carried out with saline followed by 4% depolymerized paraformaldehyde (PFA) in phosphate buffered saline (PBS). Extracted brains were postfixed in the same perfusion solution for two nights at 4 °C. The brains were rinsed in 0.1 M phosphate buffer (PB; 3 × 10 min) and 50-70 μm-thick coronal sections were obtained with a vibrating blade microtome (Leica VT1000 S).

Free-floating immunofluorescence labeling was performed as described by Unal et al., 2015. Briefly, individual sections were transferred to glass vials and rinsed (3 × 10 min) in PBS containing 0.3% Triton X-100 (PBS-TX) at room temperature (RT). They were immersed in a blocking solution containing 20% Normal Horse Serum (NHS) in PBS-TX for 1 h at RT. The sections were then incubated in primary antibody solutions in PBS-TX with 1% NHS for 48 h at 4 °C. Following primary antibody incubation, the sections were rinsed in PBS-TX (3 × 10 min) and transferred to secondary antibody solutions (PBS-TX with 1% NHS) for 4 h at RT.

We used the following primary antibodies (dilution, code, company): rabbit anti-parvalbumin (PV; 1:2000, ab11427, Abcam), goat anti-choline acetyltransferase (ChAT; 1:500, AB144P, Merck-Millipore), goat anti-ChAT (1:350, ab254118, Abcam), rabbit anti-Leu-enkephalin (Leu-enk; 1:1000, ab22619, Abcam) and rabbit anti-GABA (1:1000, A2052, Sigma). The secondary antibodies were donkey anti-rabbit Alexa Fluor 488 (1:250, ab150073, Abcam) and donkey anti-goat DyLight 650 (1:1000, ab96938, Abcam).

In addition to immunohistochemistry for Leu-enk (Fig. 2C), a standard cresyl violet or DAPI (1:2000, D3571, ThermoFisher) staining was utilized in a subset of sections to delineate the borders of basal forebrain nuclei. For DAPI staining, sections were incubated in glass vials for 15 minutes and rinsed in PBS-TX (3 × 10 min) before mounting on glass slides.

**Figure 2.**
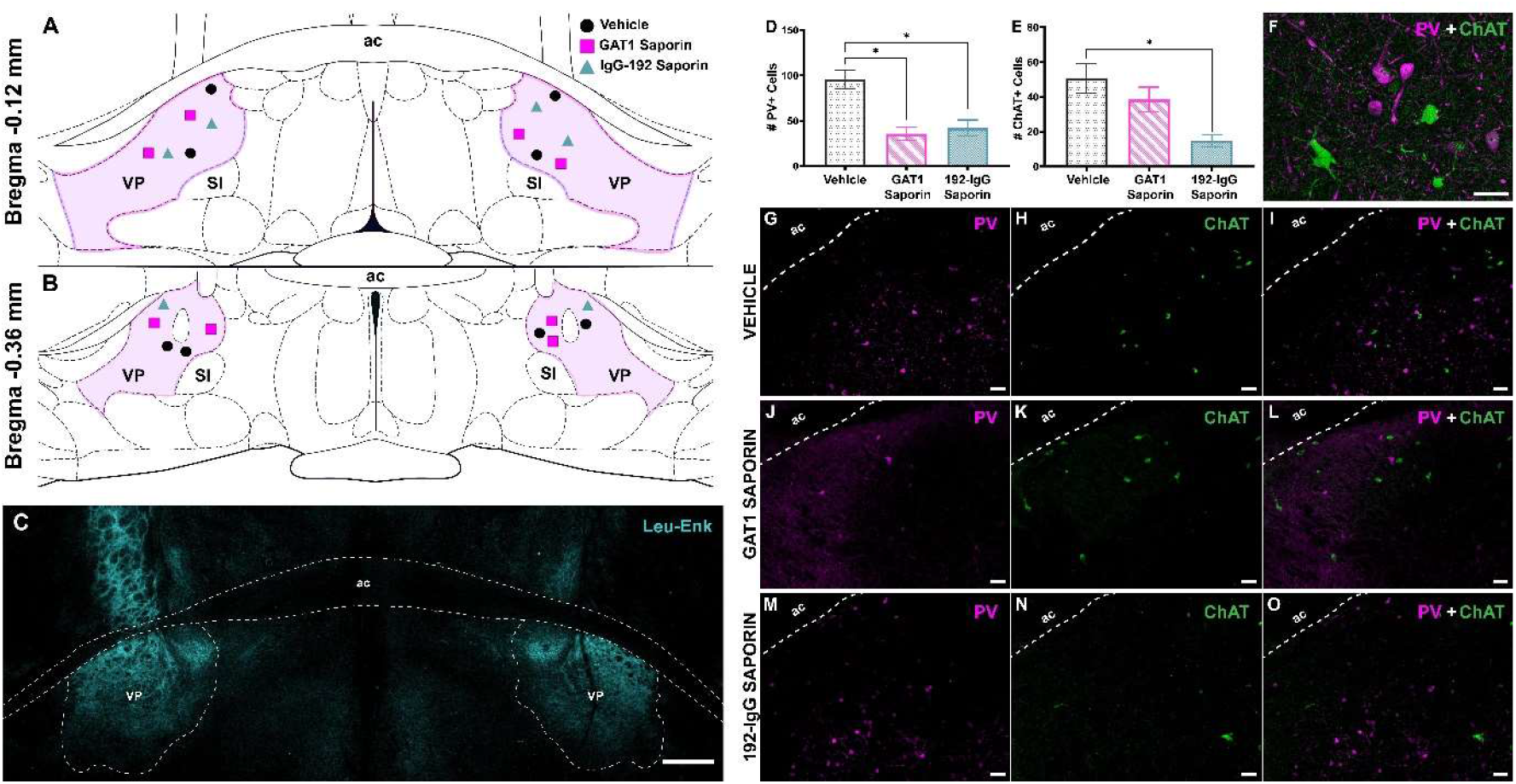
Histological and immunohistochemical verification of GAT1-Saporin and 192-IgG-Saporin lesions. (A-B) Schematic depiction of coronal sections demonstrating the centers of injection sites (n = 11 animals). (C) Fluorescent immunohistochemistry against Leu-enkephalin showing dense immunopositive fibers in the VP and globus pallidus with relatively sparse labeling in the bed nucleus of stria terminalis and medial preoptic nuclei. Scale bar: 500 μm. (D) Average number of PV+ VP cells per section (E) Average number of ChAT+ VP cells per section (F) Maximum intensity projection of a high-power confocal Z-stack in the VP of a vehicle animal showing specific PV and ChAT labeling. (G-O) Representative micrographs depicting PV and ChAT immunoreactivity in the VP of a control (G-I), GAT1-Saporin (J-L) and 192-IgG-Saporin (M-O) animal. Scale bars (F-O): 50 μm. Error bars show SEM. ac: anterior commissure. * =*p* < 0.05.

Immunofluorescence analyses were carried out with 12 randomly chosen brains (n = 4 for each group) to determine the specificity and degree of neuronal loss in the VP. We quantified the number of PV and ChAT immunoreactive cell bodies within the VP in 60 randomly selected sections (n = 20 for each group) with an Olympus BX43 epifluorescence microscope. The quantifications were used to generate the average number of PV and ChAT immunopositive neurons per unit of VP volume in each brain.

### Microscopy

Coronal sections including the target VP regions were initially observed with an Olympus BX43 epifluorescence microscope to detect the injection trace and associated lesions (Fig. 2A-B). Multichannel fluorescence observations were made using this microscope or a Leica SP7 confocal microscope equipped with a 20x (Plan Fluotar, N.A. = 0.4, dry, Leica Microsystems) and a 40x (Plan Apochromat, N.A. = 1.10, water-immersion, Leica Microsystems) objective lens. The step size was set at half the optical section thickness for acquisition of z-stacks. Brightness and contrast adjustments of the micrographs were performed uniformly in FIJI (Schindelin et al., 2012). No nonlinear or selective adjustment were made in the acquired images.

### Statistical Analysis

All data was tested for normality and sphericity before hypothesis testing. Between-group analyses for normally distributed data were performed by using one-way ANOVAs. Statistically significant main effects were followed by post-hoc comparisons between the vehicle and immunotoxin groups using Fisher’s LSD test. Behavioral tests with multiple trials or days were analyzed with two-way mixed ANOVA designs. When the sphericity assumption was violated, Greenhouse-Geisser correction was used and the corrected statistics were reported. All statistical analyses were performed with two-tailed tests at α = 0.05 using GraphPad Prism version 9.0.0 (GraphPad Software, San Diego, California).

## Results

### Selective Lesioning

Detailed histological analysis and fluorescent immunohistochemistry were performed on each brain in order to verify the boundaries and effectiveness of immunotoxin injections. Estimated center of each injection site was observed within the boundaries of the VP in both hemispheres (Fig. 2A-B), as delineated by densely labeled Leu-enkephalin-positive processes (Fig. 2C).

Fluorescent immunohistochemistry for PV and ChAT produced specific labeling of the two neuronal populations (Fig. 2F). Immunotoxin injections significantly altered both the PV-immunopositive (PV+) (*F*(2,9) = 13.930, *p* = 0.0018, one-way ANOVA; Fig. 2D) and ChAT-immunopositive (ChAT+) neurons (*F*(2,9) = 7.382, *p* = 0.0127, one-way ANOVA; Fig. 2E) in the VP. Compared to the PV (95.54 ± 20.14) and ChAT (50.46 ± 17.03) immunoreactivity in the vehicle group, GAT1-Saporin injections led to a significant loss of PV+ neurons (35.71 ± 14.89; *t*(9) = 4.801, *p* = 0.0010, Fisher’s LSD, Fig. 2D) while sparing ChAT+ cells (38.46 ± 14.22; *t*(9) = 1.264, *p* = 0.2380, Fisher’s LSD, Fig. 2E) in the VP. 192-IgG-Saporin injections caused a significant reduction in the number of both ChAT+ (14.63 ± 6.96; *t*(9) = 3.775, *p* = 0.0044, Fisher’s LSD, Fig. 2E) and PV+ (41.96 ± 17.45; *t*(9) = 4.300, *p* = 0.0020, Fisher’s LSD, Fig. 2D) neurons (Fig. 2G-O) as previously reported (Torres et al., 1994).

An overall quantification revealed that PV+ neurons in the VP were reduced by 63% following GAT1-Saporin injections, while192-IgG-Saporin led to a 71% reduction in the number of ChAT+ neurons. Similar ratios were found by earlier studies (see Dwyer et al., 2007; Pang et al., 2001, 2011; Roland et al., 2014; Yoder & Pang, 2005).

### Behavioral Despair

Immunotoxin administration into the VP produced a significant main effect on overall immobility during the second day of the FST (*F*(2, 21) = 7.564, *p* = 0.0034, one-way ANOVA; Fig. 3A). Both GAT1-Saporin (64.38 ± 40.19 s, *t*(21) = 3.310, *p* = 0.0053, Fisher’s LSD) and 192-IgG-Saporin (55.63 ± 31.47 s, *t*(21) = 3.578, *p* = 0.0018, Fisher’s LSD) animals displayed less immobility compared to the vehicle group (122.5 ± 39.82 s) during FST-2. In contrast, swimming patterns during the test phase of the FST did not change between the groups (*F*(2, 21) = 0.665, *p* = 0.5247, one-way ANOVA; Fig. 3B). Animals in the vehicle (34.69 ± 19.84 s), GAT1-Saporin (30.63 ± 15.45 s) and 192-IgG-Saporin (43.44 ± 30.24 s) groups showed similar levels of swimming.

**Figure 3.**
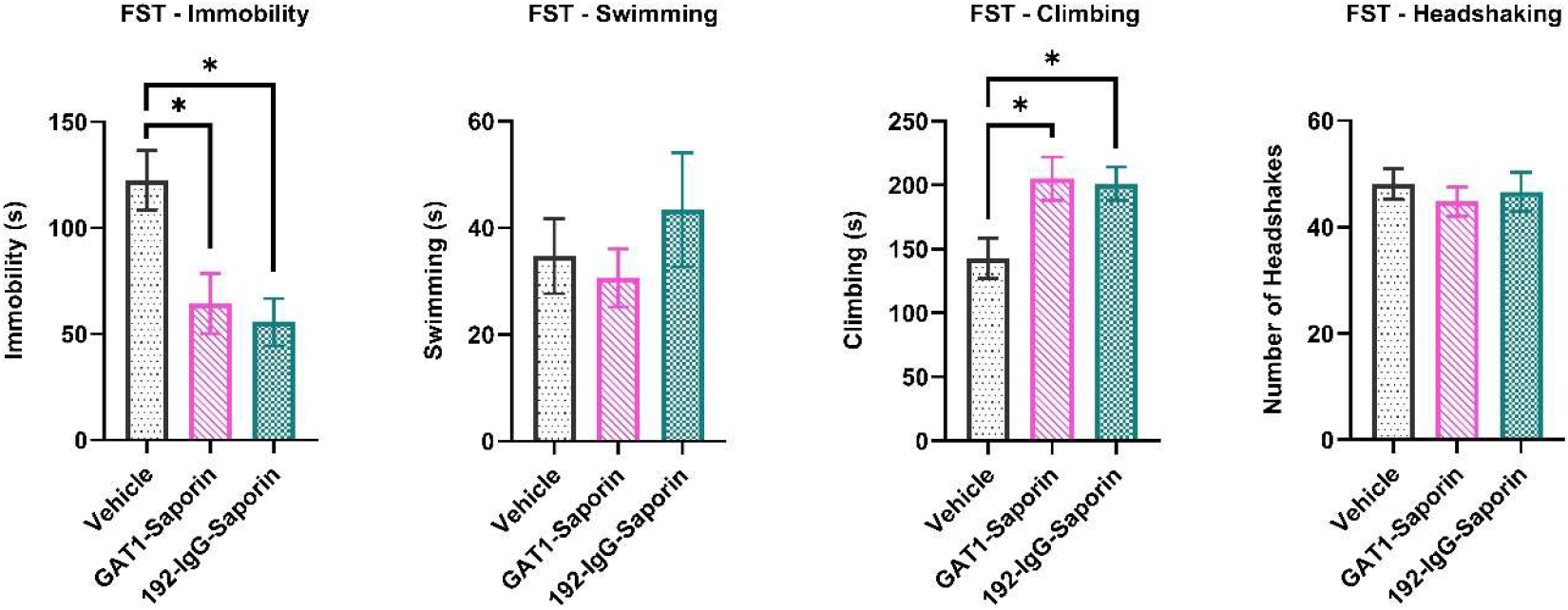
Behavioral despair analyses from the 5-minute test session of the FST. (A-C) Average durations of immobility (A), swimming (B) and climbing/struggling (C). (D) Average number of headshakes. Error bars show SEM. * =*p* < 0.05.

Saporin injections increased the duration of climbing behavior (*F*(2, 21) = 5.113, *p* = 0.0155, one-way ANOVA; Fig. 3C) during FST-2. Animals that received intra-VP GAT1-Saporin (205 ± 48.05 s, *t*(21) = 2.858, *p* = 0.0094, Fisher’s LSD) and 192-IgG-Saporin (200.9 ± 37.20 s, *t*(21) = 2.671, *p* = 0.0143, Fisher’s LSD) injections displayed longer periods of climbing compared to the vehicle group (142.8 ± 44.59 s). There were no group-level differences in headshaking (*F*(2, 21) = 0.283, *p* = 0.7561, one-way ANOVA; Fig. 3D) or diving (*F*(2, 21) = 1.382, *p* = 0.2730, one-way ANOVA) in the FST-2.

### Locomotor Activity and Anxiety

There was no group-level difference in total distance travelled (*F*(2, 20) = 1.249, *p* = 0.3082, one-way ANOVA; Fig. 4A) or the average speed of locomotion in the OFT (*F*(2, 20) = 1.535, *p* = 0.2398, one-way ANOVA), indicating that the antidepressant-like effect observed in the FST was not due to alterations in general locomotor activity. One animal from the 192-IgG-Saporin group showed excessive freezing in the OFT and did not leave the starting position at the center of the maze (time spent in the center = 300 s vs. group mean = 27.88 ± 24.19 s, z = 11.2513). This outlier was excluded from the OFT analyses.

**Figure 4.**
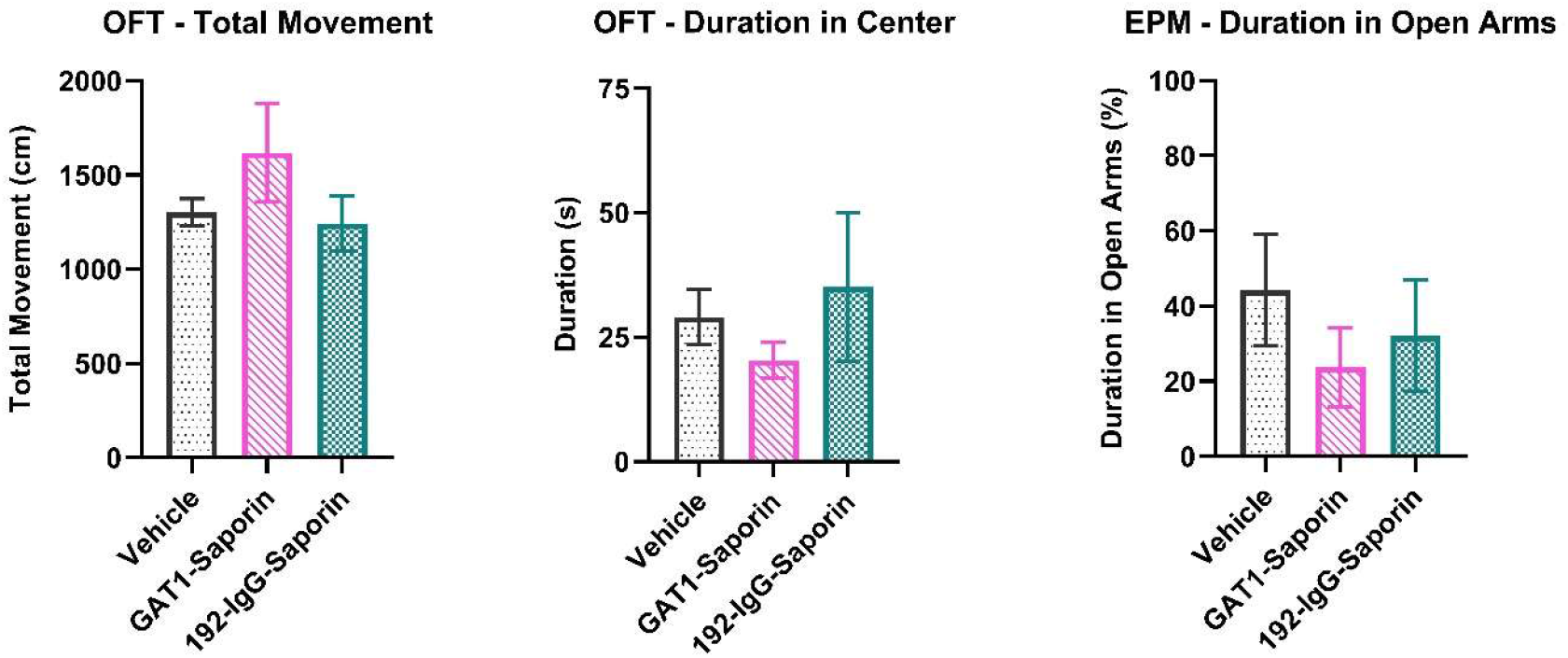
Assessment of locomotor activity and anxiety-like behavior in the OFT and EPM. (A) Total movement/distance travelled (cm) during the OFT. (B) Average time (s) spent in the central zone of the OFT. (C) Percentage of time spent in the open arms of the EPM. Error bars show SEM. * =*p* < 0.05.

There was no difference in anxiety-like behavior observed in the OFT, as assessed by comparing the overall time spent in the central portion of the maze of each group (*F*(2, 20) = 0.6952, *p* = 0.5106, one-way ANOVA; Fig. 4B). Similarly, percentage of time spent in the open arms of the EPM were not different between the groups (*F*(2, 21) = 0.5880, *p* = 0.5643, one-way ANOVA; Fig. 4C). The number of crossings to open arms did not differ following immunotoxin injections (*F*(2, 21) = 1.528, *p* = 0.2401, one-way ANOVA). All animals showed similar locomotor activity patterns in the OFT, which were dominated by thigmotaxis.

### Spatial Learning and Memory

MWM analyses revealed a significant decrease in escape latency after four days of training (F(1.782, 37.43) = 22.78, *p* < 0.0001, 3×4 two-way mixed ANOVA; Fig. 5A), indicating that spatial learning was achieved by the final day of training. We also found a significant main effect of manipulation on the overall time spent to locate the platform (*F*(2, 21) = 5.437, *p* = 0.0125, 3×4 two-way mixed ANOVA). The significant difference based on the immunotoxin injections emerged mainly from the impaired spatial navigation performance of the 192-IgG-Saporin animals. 192-IgG-Saporin animals were slower to find the platform on the second (*t*(21) = 3.550, *p* = 0.0032, Fisher’s LSD), third (*t*(21) = 3.314, *p* = 0.0079, Fisher’s LSD), and the fourth (*t*(21) = 2.579, *p* = 0.0291, Fisher’s LSD) days of training (Fig. 5A) when compared to the vehicle group.

**Figure 5.**
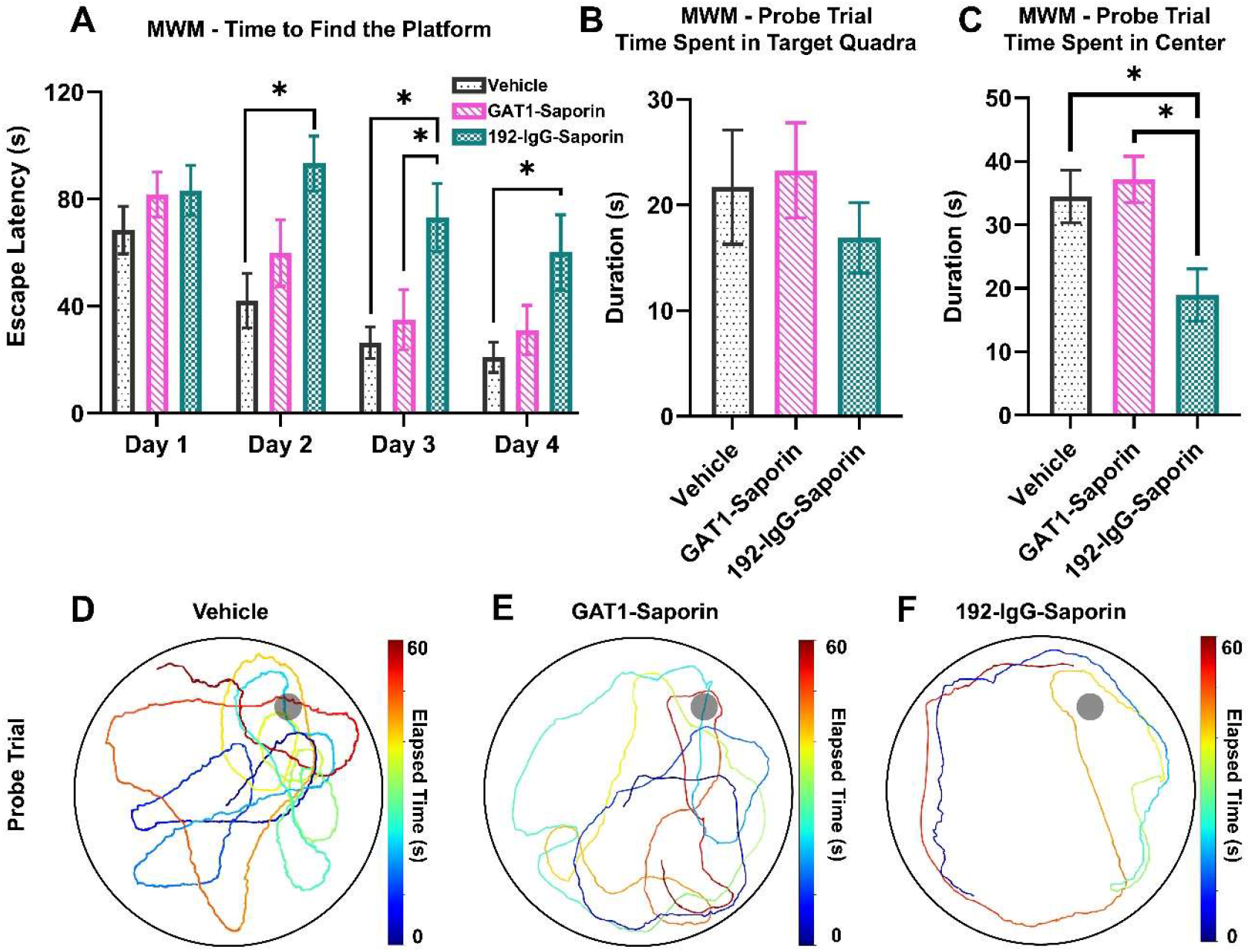
Spatial learning assessed in the MWM. (A) Average escape latency (s) scores per training day. (B) Average time (s) spent in the target (platform) quadrant during the probe trial. (C) Average time (s) spent in the central area of the MWM during the probe trial. (D-F) Representative swimming trajectories from each group during the probe trial. The trajectories were created using a custom DLC implementation. Error bars show SEM. * =*p* < 0.05.

During the probe trial, the three groups showed no difference in time spent in the target quadrant (*F*(2, 21) = 0.5489, *p* = 0.5857, one-way ANOVA; Fig. 5B) or the average number of platform crossings (*F*(2, 21) = 1.812, *p* = 0.1879, one-way ANOVA; Fig. 5B). The average distance traveled (*F*(2, 21) = 0.2887, *p* = 0.7522, one-way ANOVA) or the average speed of the animals (*F*(2, 21) = 1.62, *p* = 0.2213, one-way ANOVA) also did not differ between the groups. However, the movement patterns of the groups were significantly different in the probe trial (*F*(2, 21) = 6.064, *p* = 0.0083, one-way ANOVA; Fig. 5C). 192-IgG-Saporin animals (18.96 ± 11.65 s) spent significantly less time in the central zone of the MWM as compared to the vehicle (34.47 ± 11.73 s, *t*(21) = 2.750, *p* = 0.120, Fisher’s LSD) and GAT1-saporin (37.14 ± 10.39 s, *t*(21) = 3.225, *p* = 0.0041, Fisher’s LSD) injected groups during the probe trial. While vehicle injections (Fig. 5D) and GAT1 lesions (Fig. 5E) produced a similar, more distributed swimming pattern, cholinergic lesions lead to predominant thigmotaxis in the probe trial (Fig. 5F). The groups did not differ in the average distance traveled (*F*(2, 21) = 0.2887, *p* = 0.7522, one-way ANOVA) or average speed (*F*(2, 21) = 1.622, *p* = 0.2213, one-way ANOVA) in the probe trial.

### Natural and Conditioned Fear Response

Pavlovian fear conditioning to an auditory CS was followed by extinction sessions in the same (context A) or a different apparatus (context B), respectively three and seven days after conditioning (Fig. 6A). Immunotoxin injections into the VP altered both the unconditioned and conditioned fear responses observed during conditioning. US presentations during fear conditioning led to increased darting behavior in GAT1-Saporin (4.00 ± 2.00; *t*(21) = 2.548, p = 0.0187, Fisher’s LSD) and 192-IgG-Saporin (4.50 ± 1.93; *t*(21) = 3.114, *p* = 0.0052, Fisher’s LSD) animals compared to the vehicle group (1.75 ± 1.28; *F*(2, 21) = 5.504, *p* = 0.0120, one-way ANOVA; Fig. 6B). The animals also showed an altered frequency of jumping behaviors in response to the US based on their experimental groups (*F*(2, 21) = 13.45, *p* = 0.0002, one-way ANOVA, Fig. 6C). GAT1-Saporin animals displayed a significantly higher frequency of jumping in response to footshocks (2.13 ± 0.64; *t*(21) = 2.954, *p* = 0.0076, Fisher’s LSD) compared to the vehicle group (1.13 ± 0.83). We found that 192-IgG-Saporin animals (0.38 ± 0.52), on average, jumped significantly less during conditioning compared to both the vehicle (*t*(21) = 2.216, *p* = 0.0379, Fisher’s LSD) and the GAT1-saporin injected animals (*t*(21) = 5.170, *p* < 0.0001, Fisher’s LSD).

**Figure 6.**
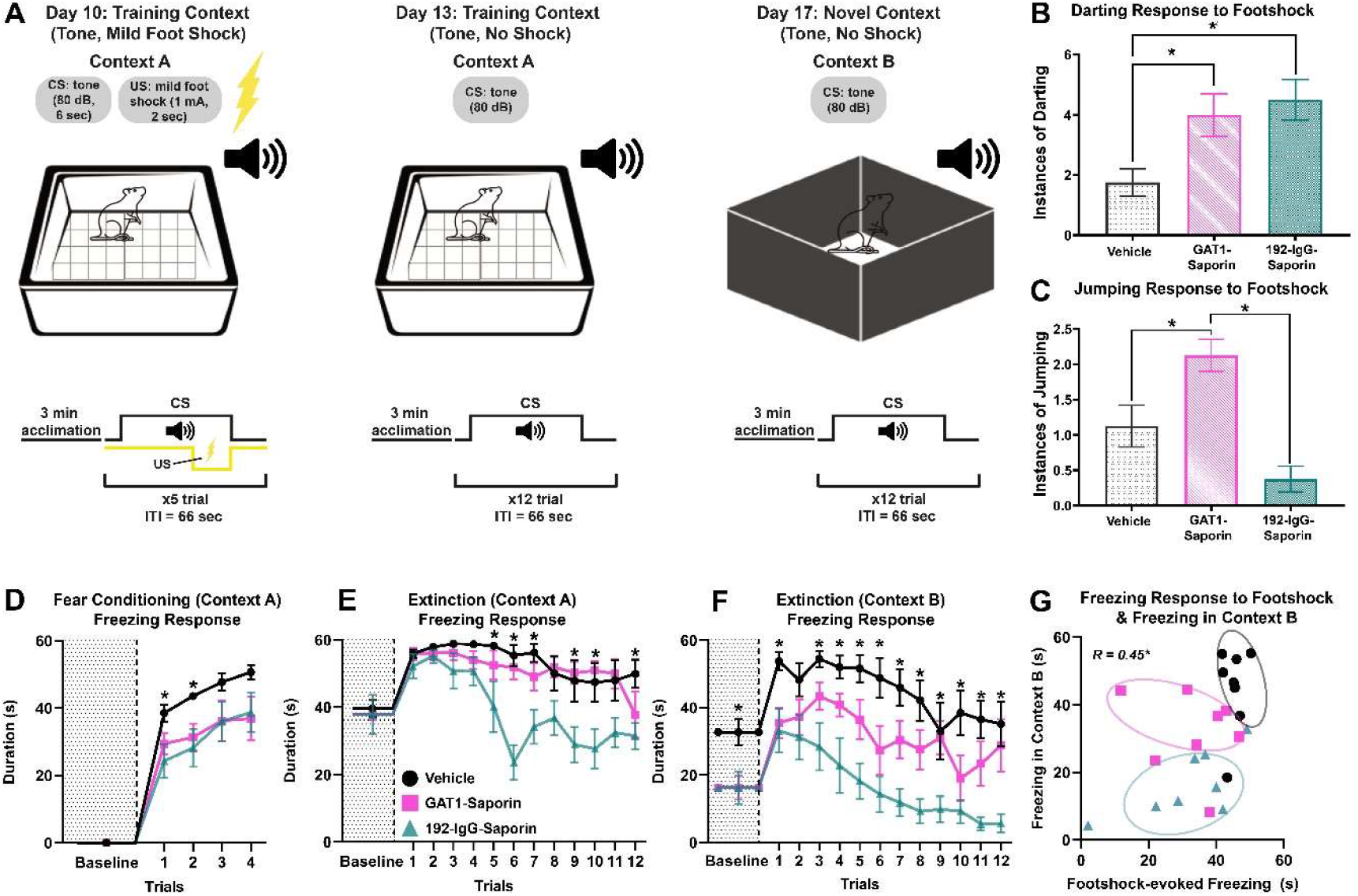
Auditory fear conditioning. (A) Schematic depiction of conditioning and extinction sessions. (B) Average instances of darting in response to footshock. (C) Average instances of jumping in response to footshock. (D) Baseline and average conditioned freezing per trial (context A). (E) Baseline and average conditioned freezing per trial during extinction in the conditioning apparatus (context A). (F) Baseline and average conditioned freezing per trial during extinction in a novel apparatus (context B). (G) Scatter plot of the correlation between average freezing in response to footshock and auditory cued freezing in context B. The distinct clusters formed by the three groups are encircled. Error bars show SEM. * =*p* < 0.05.

Fear conditioning and extinction sessions started with a 3-min acclimation period during which baseline freezing was recorded. None of the animals exhibited freezing before conditioning (Fig. 6D). Freezing emerged with the first CS-US pairing and persisted through conditioning for all groups (*F*(2.217, 46.57) = 149.3, *p* < 0.0001, 3×4 two-way mixed ANOVA; Fig 6D). We observed a group-level difference resulting from the first two US-CS pairings (*F*(2, 21) = 3.553, *p* = 0.0469, two-way mixed ANOVA). Vehicle animals displayed heightened freezing (38.50 ± 7.13) compared to the GAT1-Saporin (29.38 ± 8.81; *t*(13.42) = 2.276, *p* = 0.0398, Fisher’s LSD) and the 192-IgG-Saporin group (24.31±13.92; *t*(10.44) = 2.566, *p* = 0.0272, Fisher’s LSD). The difference persisted in the second pairing, as control animals further increased their freezing response (43.56 ± 2.72) compared to the GAT1-Saporin (31.38 ± 11.44; *t*(7.79) = 2.932, *p* = 0.0195, Fisher’s LSD) and 192-IgG-Saporin group (28.20 ± 16.07; *t*(7.40) = 2.666, *p* = 0.0306, Fisher’s LSD; Fig. 6D).

Neither GAT1-Saporin nor 192-IgG-Saporin injections had any significant effect on freezing behavior of the groups during the 3-min baseline period in the extinction session conducted in Context A (*F*(2, 21) = 0.0746, *p* = 0.9284, one-way ANOVA; Fig. 6E). Immunotoxin lesions led to altered freezing in the extinction trials in Context A (*F*(2, 21) = 8.124, *p* = 0.0024, 3×12 two-way mixed ANOVA) with a significant trial-lesion interaction (*F*(22, 231) = 2.746, *p* < 0.0001, 3×12 two-way mixed ANOVA; Fig. 6E). 192-IgG-Saporin animals showed less freezing (40.21 ± 21.20) compared to the vehicle (58.24 ± 2.62; *t*(7.21) = 2.387, *p* = 0.0473, Fisher’s LSD) group starting with the fifth trial. With the following (sixth) trial, 192-IgG-Saporin animals (23.65 ± 14.59) also showed less freezing compared to the GAT1-Saporin (51.84 ± 10.39; *t*(12.65) = 4.453, *p* = 0.0007, Fisher’s LSD) group in addition to the vehicle group (55.36 ± 9.06; *t*(11.70) = 5.225, *p* = 0.0002, Fisher’s LSD).

Immunotoxin injections significantly altered freezing behavior during the 3-min baseline period in the extinction test conducted in Context B (*F*(2, 21) = 5.319, *p* = 0.0135, one-way ANOVA; Fig. 6F). Both the GAT1-Saporin (16.29 ± 9.94; *t*(21) = 2.822, *p* = 0.0102, Fisher’s LSD) and 192-IgG-Saporin (16.25 ± 13.61; *t*(21) = 2.828, *p* = 0.0101, Fisher’s LSD) groups showed significantly less freezing during the baseline measurement in Context B compared to the vehicle group (32.71 ± 11.05). Extinction in Context B also revealed a significant effect of experimental manipulation (*F*(2, 21) = 12.32, *p* = 0.0003, 3×12 two-way mixed ANOVA; Fig. 6F). As in Context A, cholinergic lesions significantly and consistently decreased freezing compared to the vehicle group in the majority of trials, this time starting with the first trial (192-IgG-Saporin: 33.16 ± 18.56; Vehicle: 53.84 ± 7.34; *t*(9.14) = 2.930, *p* = 0.0165, Fisher’s LSD). In this novel context, however, GAT1-Saporin animals (36.81 ± 14.24) also showed significantly less freezing on average in the first 6 trials when compared to the vehicle group (51.52 ± 11.03; *t*(14) = 2.467, *p* = 0.0135, between-groups t-test). This effect did not persist in the second half of the extinction trials in context B (*t*(14) = 1.713, *p* = 0.0544, between-groups t-test). Thus, we observed impaired conditioned fear memory in the 192-IgG-Saporin group, and to a lesser extent in GAT1-Saporin animals when context-related information was missing.

Exploratory correlation analyses revealed a significant correlation between the average duration of freezing per trial observed in the conditioning and Context B extinction sessions (*r*(22) = 0.4485, *p* = 0.0279). The three groups formed distinct clusters in this exploratory space (Fig. 6G). The vehicle group showed high levels of freezing with relatively low variability both in conditioning and Context B extinction. The 192-IgG-Saporin group was characterized by significantly lower levels of cued freezing, while the freezing scores of GAT1-Saporin animals were distributed between the other two groups (Fig. 6G).

## Discussion

We showed that selective GABAergic or cholinergic VP lesions led to altered behavior in different affective and cognitive processes, both of which were marked by an overall increase in the frequency of active responses to aversive stimuli. A substantial antidepressant-like effect and higher levels of struggling behavior were observed for both groups in the FST following saporin injection. In terms of cognitive processioning, 192-IgG-Saporin, but not GAT1-Saporin, lesions caused a decline of spatial memory performance as assessed in the MWM.

Ventral pallidum is a core component of the reward circuitry of the mammalian brain, with well-known functions in motivated and appetitive behaviors (Ottenheimer et al., 2020; Root et al., 2015). In contrast, its role in aversive processing has only recently attracted scientific attention (Farrell et al., 2021; Moaddab et al., 2021; Saga et al., 2017; Stephenson-Jones et al., 2020). We investigated this understudied system by selectively targeting the GABAergic and cholinergic populations of the VP, and revealed that both neuronal groups contribute to behavioral despair and conditioned fear responses by suppressing active coping. Climbing, or struggling, in the FST chamber as well as darting and jumping responses to footshock in fear conditioning constitute different forms of active coping. Our observations on these active coping mechanisms corroborate existing electrophysiological (Moaddab et al., 2021) and behavioral evidence (Saga et al., 2017), pointing to a crucial role for VP GABAergic neurons in encoding aversive stimuli and producing adaptive responses. They also shed light on an earlier study observing an increase in digging and forepaw treading following bicuculline, a GABA-A receptor antagonist, injections into the VP (Smith & Berridge, 2005). These species-specific behaviors resemble the active coping that emerged under controlled behavioral paradigms of this study.

### Depressive-Like Behavior

The VP has reciprocal connections with several brain regions that are directly implicated in the pathophysiology of clinical depression as well as depressive-like behaviors in rodents. These include the nucleus accumbens, lateral habenula, ventral tegmental area, amygdala, and prefrontal cortex (Gielow & Zaborszky, 2017; Groenewegen et al., 1993; Haber et al., 1985; Hakan et al., 1992; Heimer et al., 1991; Phillipson & Griffiths, 1985; Root et al., 2015; Taylor et al., 2014; Zahm et al., 1996). Despite these connections and partly due to its regulatory function in appetitive behaviors, the role of VP in affective disorders has largely been neglected in neuroscience research (Root et al., 2015). Here, we found that selective deletion of GABAergic or cholinergic VP neurons produced potent antidepressant-like effects in the FST.

It is important to note the FST, the most common behavioral rodent test of antidepressant efficacy, has been criticized for its construct validity. The main measure of this test is immobility, which corresponds to the rodent endophenotype of psychomotor retardation, a symptom required for clinical diagnosis of major depressive disorder (American Psychiatric Association, 2013; Unal & Canbeyli, 2019). In the current study, both lesion groups had decreased immobility in the test phase compared to the vehicle animals that displayed behavioral despair. In line with this, we found a significant increase in active climbing behavior or struggling. The FST analyses did not arise from different levels of metabolic activity between the groups as the OFT revealed similar locomotor activity levels for all animals.

An earlier study investigated the role of VP GABAergic neurons in depressive-like behavior by enhancing their function (Skirzewski et al., 2011). They showed that heightened VP GABAergic tone led to increased immobility during the test phase of the FST, while intra-VP bilateral bicuculline injections into the VP produced ameliorative effect. A later work demonstrated that two parallel circuits originating from different PV+ subpopulations of the VP target the lateral habenula or VTA and differentially contribute to different depressive-like rodent phenotypes (Knowland et al., 2017). The VP was suggested to contribute to stress-induced depressive-like behavior by suppressing dopaminergic activity through its connections with the basolateral amygdala (BLA) (Chang & Grace, 2014). In contrast to a recent conflicting report stating that optogenetic stimulation of VP GABAergic neurons does not influence behavioral despair (Li et al., 2020), our results provide the first direct functional evidence for the role of the VP GABAergic neurons in behavioral despair.

Furthermore, we showed that selective lesioning of cholinergic neurons in the VP also leads to antidepressant-like effects. The role of the cholinergic system in regulating depressive symptoms has generally been tested by pharmacological manipulation in the basal forebrain or their target structures (Dulawa & Janowsky, 2019). Systemic injections of acetylcholinesterase inhibitors into the VTA or hippocampus (Addy et al., 2015; Mineur et al., 2013; Small et al., 2016) as well as intra-VTA, or intra-NAc injections of cholinergic receptor agonists (Andreasen & Redrobe, 2009; Chau et al., 2001; Haj-Mirzaian et al., 2015a; Small et al., 2016) produce depression-like outcomes. In contrast, muscarinic or nicotinic antagonists lead to antidepressant-like effects (Aboul-Fotouh, 2015; Addy et al., 2015; Chau et al., 2001; Ghosal et al., 2018; Haj-Mirzaian et al., 2015b; Navarria et al., 2015). Here, we showed that the antidepressant effects of suppressing or silencing the cholinergic system extend to the VP. This indicates that basal forebrain cholinergic neurons uniformly contribute to depression-like behavior. It is, however, likely that cholinergic neurons that localize in different subnuclei and target different limbic regions contribute to different depressive symptoms.

### Conditioned Fear Memory and Active Coping

Baseline recordings prior to auditory fear conditioning showed no freezing in any of the animals before the presentation of the footshock (US). Interestingly, the vehicle group displayed significantly higher freezing to the first CS-US pairing as compared to the immunotoxin groups (Trial 1 in Fig. 6D). As this was the very first presentation of the footshock, we conclude that selective lesions of the VP GABAergic or cholinergic neurons altered the unconditioned, or natural, fear response to an aversive stimulus. This was accompanied by enhanced darting responses in both groups and increased shock-triggered jumping in the GAT1-Saporin animals. We can conclude that the extra freezing displayed by the vehicle animals was replaced by active coping in GABAergic or cholinergic lesions. Diminished unconditioned freezing observed in the experimental groups did not persist after conditioning as the baseline freezing of all three groups were leveled before the extinction session in the same context (Baseline in Fig. 6E). Cholinergic, but not GABAergic lesions impaired context-dependent fear memory during this session in the same context. However, it is important to note that the decrease in freezing started with the fifth trial. For this reason, it is also possible to state that VP cholinergic lesions do not impair acquired fear memory, but instead lead to better extinction learning (Knox, 2016; Wilson & Fadel, 2017). This is a less likely explanation as the 192-IgG-Saporin group in this study was characterized by deficits in spatial learning assessed in the MWM.

Cholinergic inputs increase signal-to-noise ratio in the BLA (Unal et al., 2015). In previous research, systemic application of cholinergic antagonists lead to decreased freezing during fear acquisition (Anagnostaras et al., 1999; Figueredo et al., 2008; Fornari et al., 2000; Soares et al., 2006). They cause memory deficits in context conditioning (Anagnostaras et al., 1995; Figueredo et al., 2008; Fornari et al., 2000), cued fear conditioning (Young et al., 1995), or both context and cued conditioning when tested together (Anagnostaras et al., 1999; Feiro & Gould, 2005; Rudy, 1996). High doses of scopolamine (Anagnostaras et al., 1999) or inhibition of basal forebrain cholinergic terminals in the BLA (Jiang et al., 2016) eliminated freezing during fear acquisition without altering pain sensitivity-related measures such as flinching, jumping or vocalization. A recent chemogenetic study showed that inhibition of VP GAD1-Cre neurons affected the motivation to avoid an aversive stimulus, but did not alter shock reactivity (Farrell et al., 2021). Similarly, we showed that selective cholinergic lesions led to enhanced darting behavior during fear acquisition, but not did not increase the number of shock-driven jumping.

Baseline recordings of the extinction session in a novel context indicate that the vehicle group could generalize the context information of the cued conditioning (Baseline in Fig. 6F). This was followed by both immunotoxin groups showing diminished conditioned freezing in a novel context. Unlike 192-IgG-Saporin animals, the impairment of the GABAergic lesions was restricted to the novel context. The differences observed in active coping and conditioned freezing altogether suggest a differential role for the GABAergic and cholinergic VP neurons in acquired fear memory.

The revealed role of VP GABAergic neurons in behavioral despair and fear memory likely arises via their connections to the amygdaloid circuitry (Mascagni & McDonald, 2009; Mcdonald et al., 2011b; McDonald et al., 2012). GABAergic neurons of the VP are involved in the production of prepulse inhibition of acoustic startle, a BLA-dependent phenomenon (Kodsi & Swerdlow, 1995, 1997; Swerdlow et al., 1990). Bilateral muscimol injections into the VP following BLA inactivation restores prepulse inhibition to normal levels (Forcelli et al., 2012). Long-range GABAergic projections of the VP may function as a mediator for amygdala-dependent acquired behaviors.

The present study provided direct evidence for the involvement of VP in behavioral responses to aversive stimuli. We showed that the GABAergic and cholinergic neuronal populations of the VP are involved in depression-like phenotypes, unconditioned/natural fear responses and different aspects of acquired fear memory. Selective lesioning of these groups produced antidepressant effects and increased active coping responses. These findings suggest that the GABAergic and cholinergic neuronal groups of the VP may constitute separate but complementary therapeutic targets for the treatment of different mood disorders.

## Acknowledgements

This research was funded by an EMBO Installation Grant to GU. The authors thank Sena Işik and Pinar Ersoy for contributing to behavioral analyses.

## Notes

1 Conflict of interest: The authors declare no competing financial interests.

### Competing Interest Statement

The authors have declared no competing interest.

